# Association of *AKAP6* and *MIR2113* with cognitive performance in a population based sample of older adults

**DOI:** 10.1101/070409

**Authors:** Shea J. Andrews, Debjani Das, Kaarin J. Anstey, Simon Easteal

## Abstract

Genetic factors make a substantial contribution to inter-individual variability in cognitive function. A recent meta-analysis of genome-wide association studies identified two loci, *AKAP6* and *MIR2113* that are associated with general cognitive function. Here, we extend this previous research by investigating the association of *MIR2113* and *AKAP6* with baseline and longitudinal nonlinear change across a broad spectrum of cognitive domains in community-based cohort of 1,570 older adults without dementia. Two SNPs, *MIR211*-rs10457441 and *AKAP6*-rs17522122 were genotyped in 1,570 non-demented older Australians of European ancestry, who were examined up to 4 times over 12 years. Linear mixed effects models were used to examine the association between *AKAP6* and *MIR2113* with cognitive performance in episodic memory, working memory, vocabulary, perceptual speed and reaction time at baseline and with linear and quadratic rates of change. *AKAP6*-rs17522122*T was associated with worse baseline performance in episodic memory, working memory, vocabulary and perceptual speed, but it was not associated with cognitive change in any domain. *MIR2113*-rs10457441*T was associated with accelerated decline in episodic memory. No other associations with baseline cognitive performance or with linear or quadratic rate or cognitive changes was observed for this SNP. These results confirm the previous finding that, *AKAP6* is associated with performance across multiple cognitive domains at baseline but not with cognitive decline, while *MIR2113* primarily affects the rate at which memory declines over time.

## Introduction

Age-associated decline is a general process that affects all cognitive domains, although it is more pronounced in domains associated with fluid cognition (i.e., that requires on-the-spot processing) than domains associated with crystalized cognition (i.e., that requires stored knowledge such as vocabulary^1^. The rate of cognitive decline in the population is, however, highly heterogeneous, with some individuals remaining relatively unimpaired across their lifecourse, while others experience a much faster rate of deterioration which can lead to cognitive impairment or dementia^2,3^. Even in the absence of dementia age-associated cognitive decline results in increased difficulty performing tasks involving memory and rapid information processing. Decline in cognitive performance is associated with poor decision-making^4^, difficulty with instrumental activities of daily living ^5,6^, poor health literacy^7^. Even when no single aspect of daily living is critically impaired, the cumulative effect of small effects in multiple domains can have a major impact on quality of life. Furthermore, a faster rate of decline is associated with dementia^2,3^ and mortality^8,9^. Identifying factors that predispose individuals to faster cognitive decline is an important step in developing intervention and treatment strategies for maintaining cognitive health.

Genetic factors contribute to the inter-individual variability observed in cognitive decline, with common genetic variants estimated to account for between 40-50% of the variability associated with general cognitive functioning in later life and 24% of the variability in lifetime cognitive change^10,11^. A recent meta-analysis of genome-wide association studies (GWAS) performed by the CHARGE consortium (n = 53,949) identified 13 SNPs in three genomic regions, *MIR2113* (n = 11), *AKAP6* (n = 1) and *APOE/TOMM40* (n = 1) associated with general cognitive function^12^. In a follow-up study of the UK Biobank (n = 112,151) *MIR2113* was associated with educational attainment and verbal-numerical reasoning, *AKAP6* was associated with verbal-numerical reasoning, reaction time and memory, while conversely *APOE/TOMM40* was not associated with any measure of cognitive function^13^.

Here, we extend this previous research by investigating the association of *MIR2113* and *AKAP6* with baseline and longitudinal nonlinear change in cognitive performance in a community-based cohort of 1,570 older adults without dementia who have undergone cognitive testing in the domains of episodic memory, working memory, verbal ability, processing speed and reaction time.

## Methods

### Participants

Participants were from the Personality and Total Health (PATH) Through Life Project, a large community survey of heath and wellbeing in adults. Participants were randomly recruited from the electoral rolls of Canberra and the neighbouring town of Queanbeyan into one of three cohorts based on age; the 20+ (20-24 years), the 40+ (40-44 years) and the 60+ (60-64 years), with individuals assessed at 4 year intervals over a period of 12 years. The background and procedures of the PATH cohort have been described in detail elsewhere ^14^. Written informed consent was obtained from all participants and approval for the study was obtained from the Human Research Ethics Committee of the Australian National University.

Data collected from the 60+ cohort was used in this study, with interviews conducted in 2001-2002 (n = 2551), 2005-2006 (n = 2222), 2009-2010 (n = 1973) and 2014-2015 (n = 1645). Individuals were excluded from analysis based on the following criteria: attendance at only 1 wave (n= 309); no genomic DNA available or missing genotype data (n = 264); *APOE* ε2/ε4 carriers (n=60, to avoid conflation of the ε2 protective and ε4 risk affect); non-European ancestry (n = 107); probable dementia at any wave (MMSE < 24; n = 82); self-reported medical history of epilepsy, brain tumours or infections, stroke and transient ischemic attacks (n = 381). Missing values in “Education” (total number of years of education, n = 129) were imputed using the ‘missForest’ package in R^15^. These exclusions left a final dataset of 1,570 individuals at baseline.

### Cognitive Assessment

All participants were assessed at baseline and at each subsequent interview for the following four cognitive abilities: perceptual speed, assessed using the Symbol Digit Modalities Test, which asks the participant to substitute as many digits for symbols as possible in 90s^16^; episodic memory, assessed using the Immediate Recall of the first trial of the California Verbal Learning Test, which involves recalling a list of 16 nouns^17^; working memory, assessed using the Digit Span Backward from the Wechsler Memory Scale, which presents participants with series of digits increasing in length at the rate of one digit per second and asks them to repeat the Digits Backwards ^18^; and vocabulary, assessed with the Spot-the-Word Test, which asks participants to choose the real words from 60 pairs of words and nonsense words^19^. Simple and choice reaction time (SRT and CRT) tasks were administrated using a hand held box with two depressible buttons, two red stimulus lights and one green ‘get ready’ light. SRT was measured using four blocks of 20 trials, in which the participant was instructed to press the right hand button (regardless of dominance) in response to the activation of one of the stimulus lights. CRT was measured using two blocks of 20 trials, in which participants were instructed to press the button corresponding to the left or right stimulus light. Mean reaction times were calculated as described previously ^20^. Raw cognitive test scores at each wave and Pearson correlations between test scores are presented in Supplementary Tables 1 & 2.

### Genotyping

The most strongly associated SNPs identified by the CHARGE consortium, *MIR211*-rs10457441, *AKAP6*-rs17522122 and *TOMM40*-rs10119, were genotyped using TaqMan OpenArray Assays as previously described^21^. For *AKAP6* and *MIR2113* an additive model was used which examines the effect of each minor allele.

*TOMM40*-rs10119 did not pass quality control in our dataset and as such was excluded from analysis. *TOMM40*-rs10119 is located in the 19q13.32 region, which is a gene-dense region of strong linkage disequilibrium and includes *APOE* and *TOMM40*. Fine mapping of the region indicates that *APOE* variation is driving observed associations with cognitive aging^22^. As such, we included the *APOE* ε2/ε3/ε4 haplotypes in our analysis as covariates. SNP data for rs429358 and rs7412 which define the APOE ε2/ε3/ε4 haplotypes, were genotyped separately using TaqMan assays as previously described^23^. APOE alleles were coded as the number of *APOE* *ε4 alleles (0,1,2). Participants with the *APOE* *ε2/ε4 allele were excluded to avoid conflation between the *APOE* *ε2 protective and *APOE* *ε4 risk affect.

All SNPs were in Hardy-Weinberg equilibrium (p < 0.05). Genotype frequencies are presented in Table 1.

**Table 1:** PATH cohort demographics

| Variable | Excluded<br>(n = 981) | Included<br>(n = 1,570) | $t/\chi^2$ | Degrees of<br>Freedom | *P* |
| --- | --- | --- | --- | --- | --- |
| Age<sup>1</sup> | 62.47 ± 1.48 | 62.54 ± 1.52 | 1.1 | 2111.22 | 0.27 |
| Education<sup>1</sup> | 13.35 ± 3.06 | 14.03 ± 2.56 | 5.81 | 1807.7 | <0.001 |
| MMSE<sup>1</sup> | 28.67 ± 2.04 | 29.37 ± 0.91 | 10.07 | 1206.95 | <0.001 |
| Male<sup>2</sup> | 513 (52.35) | 803 (51.15) | 0.4 | 1 | 0.53 |
| *MIR2113, n (%)* |  |  |  |  |  |
| T/T | 188 (26.1) | 351 (22.4) |  |  |  |
| C/T | 343 (47.6) | 805 (51.3) |  |  |  |
| C/C | 189 (26.2) | 414 (26.4) |  |  |  |
| NA<sup>3</sup> | 260 (10.2) | - |  |  |  |
| *AKAP6, n (%)* |  |  |  |  |  |
| T/T | 154 (21.5) | 342 (21.8) |  |  |  |
| G/T | 361 (50.4) | 811 (51.7) |  |  |  |
| G/G | 201 (28.1) | 417 (26.6) |  |  |  |
| NA<sup>3</sup> | 263 (10.3) | - |  |  |  |
| *APOE ε4 alleles, n (%)* |  |  |  |  |  |
| 0 | 613 (76.1) | 1183 (75.4) |  |  |  |
| 1 | 170 (21.1) | 361 (23) |  |  |  |
| 2 | 23 (2.9) | 26 (1.7) |  |  |  |
| NA<sup>3</sup> | 173 (6.7) | - |  |  |  |

<sup>1</sup>unpaired 2-tailed t-test; <sup>2</sup>Pearson's $\chi^2$ 2-tailed test; <sup>3</sup>percentage of total sample (excluded + included) with missing genotype data.

### Statistical Analysis

Statistical analysis was performed in the R 3.2.3 Computing environment ^24^. To facilitate comparisons across cognitive tests, all cognitive test results at all four waves were transformed into z-scores using the means and standard deviations at baseline. Higher scores indicate better cognitive performance. Demographic characteristics of participants excluded or included from analysis are described using means and standard deviations for continuous variables and frequency and proportions for categorical variables (Table 1). Demographic variables were compared using independent sample t-tests and χ2 tests. Linear Mixed Effects models (LMMs) adjusting for sex, years of education and *APOE* genotype, with maximum likelihood estimation, subject specific random slopes and intercepts were used to compare cognitive performance between genotypes. Longitudinal change in cognitive performance was modelled as a quadratic growth curve, where age was centred on baseline, linear rate of change (Time) was the slope at intercept and quadratic rate of change (Time^2^) was acceleration of the curve over time. Quadratic growth curves were represented as orthogonal polynomials to avoid colinearity problems and facilitate estimation of the models^25^. LMMs were estimated using the R package ‘lme4’^26^.

Statistical significance of the fixed effects were determined using a Kenward-Roger approximation for *F*-tests, where a full model, containing all fixed effects, is compared to a reduced model that excludes an individual fixed effect^27^. As this is a follow-up study to replicate the previous findings in the CHARGE consortium and UK biobank studies, a *P*-value < 0.05 was considered statistically significant. Conditional 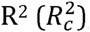, the variance explained by fixed and random effects (i.e. the entire model), and marginal 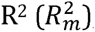, the variance explained by the fixed effects were calculated using the R package ‘MuMIn’^28–30^.

## Results

### Population Characteristics of the PATH cohort

Demographic information on PATH participants is presented in Table 1. Individuals retained in the final sample had on average, higher MMSE scores and more years of education than those excluded. Unconditional growth LMMs showed that all cognitive tests were associated with linear rate of change and, with the exception for Digits Backwards test, quadratic rate of change (supplementary table 3). Supplementary table 4 shows the results of the covariates only model.

### Main effects of *AKAP6* and *MIR2113*

Parameter estimates for the associations of *AKAP6* and *MIR2113* with cognitive performance are presented in Table 2. Figure 1 illustrates the differences in cognitive trajectories between *AKAP6* and *MIR2113* genotypes.

**Figure 1:**
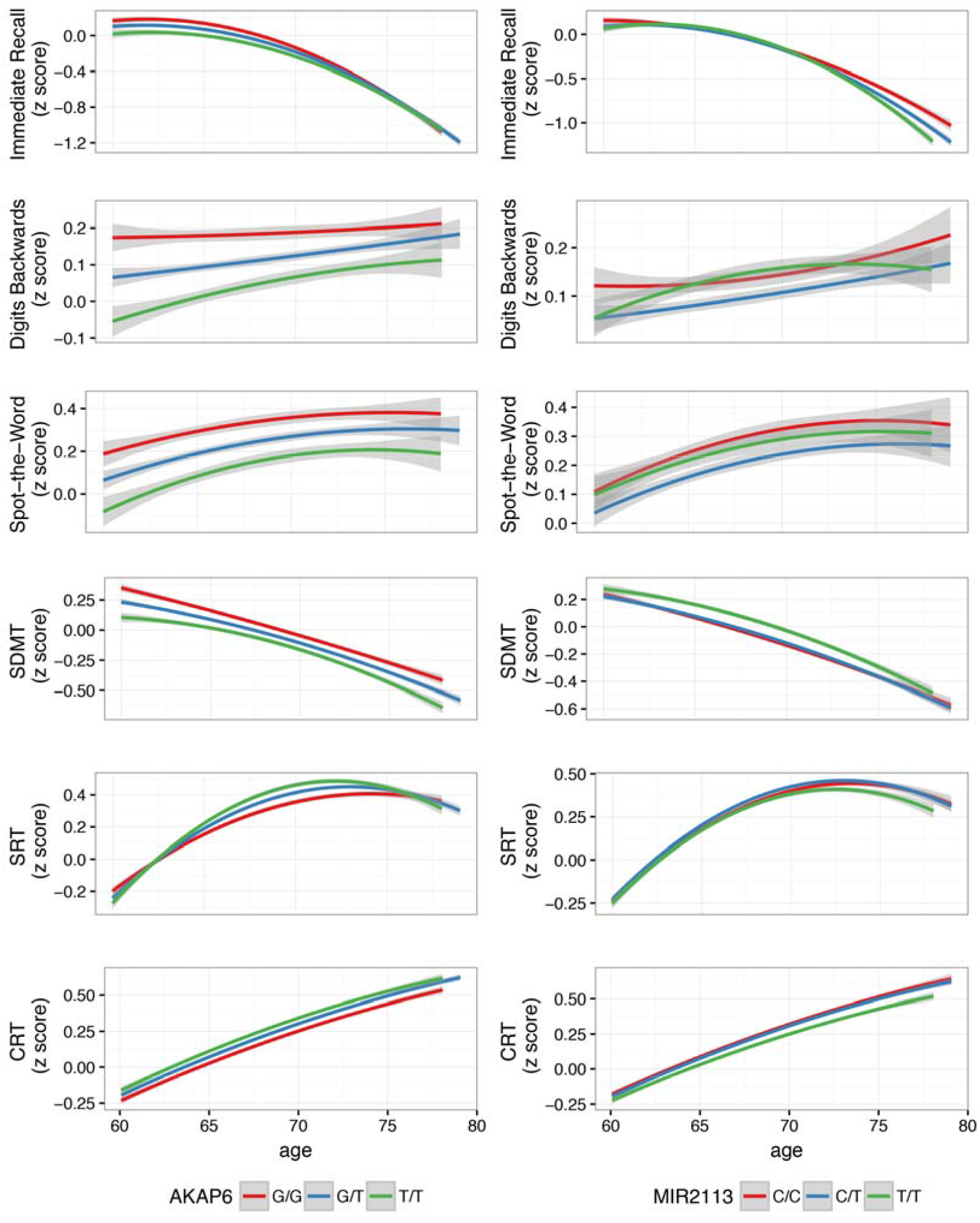
Trajectories of cognitive performance for *AKAP6* and *MIR2113* genotype

**Table 2:**
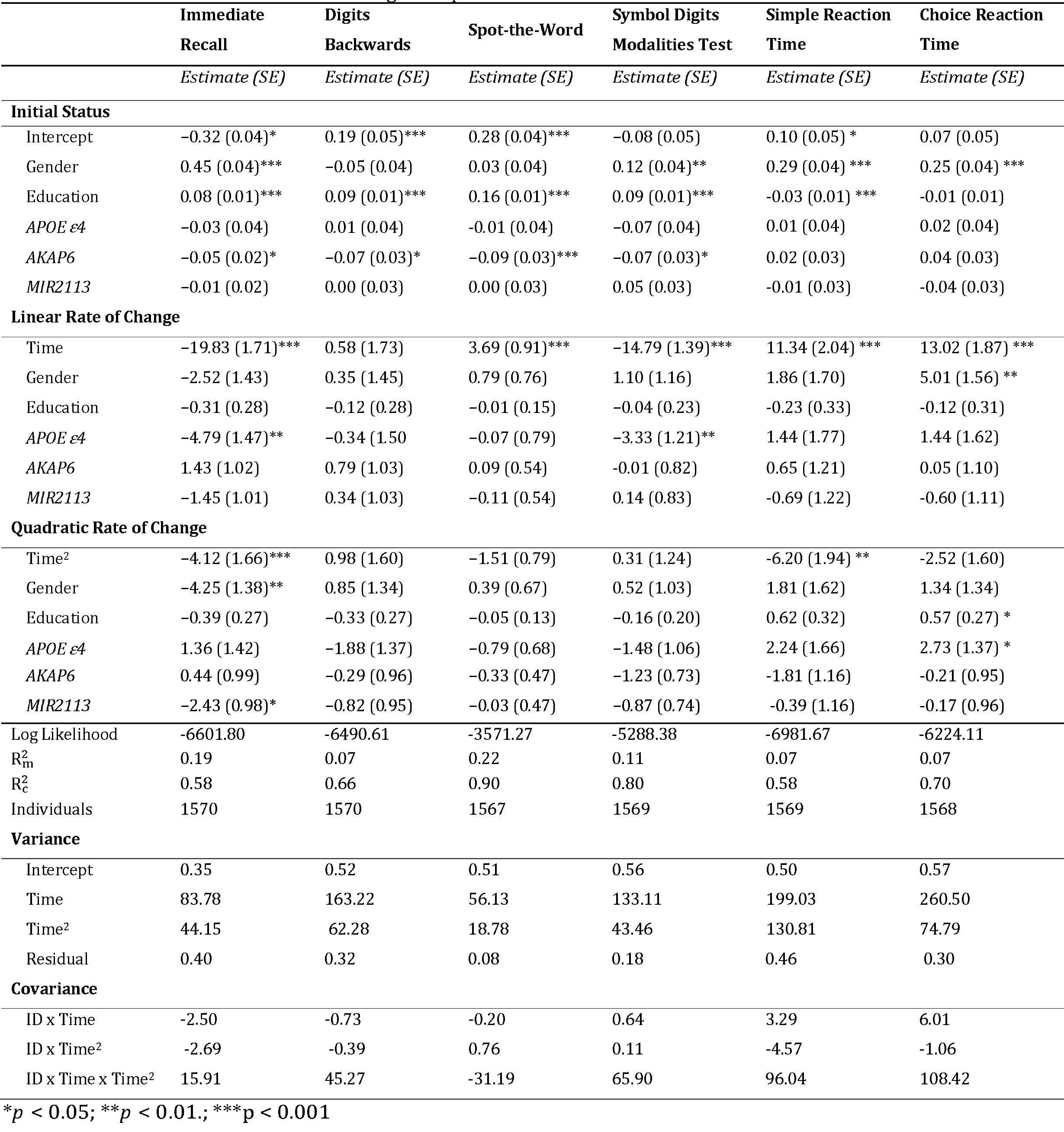
Effect of *AKAP6* and *MIR2113* on cognitive performance

|  | Immediate Recall | Digits Backwards | Spot-the-Word | Symbol Digits Modalities Test | Simple Reaction Time | Choice Reaction Time |
| --- | --- | --- | --- | --- | --- | --- |
|  | *Estimate (SE)* | *Estimate (SE)* | *Estimate (SE)* | *Estimate (SE)* | *Estimate (SE)* | *Estimate (SE)* |
| **Initial Status** |  |  |  |  |  |  |
| Intercept | -0.32 (0.04)* | 0.19 (0.05)*** | 0.28 (0.04)*** | -0.08 (0.05) | 0.10 (0.05) * | 0.07 (0.05) |
| Gender | 0.45 (0.04)*** | -0.05 (0.04) | 0.03 (0.04) | 0.12 (0.04)** | 0.29 (0.04) *** | 0.25 (0.04) *** |
| Education | 0.08 (0.01)*** | 0.09 (0.01)*** | 0.16 (0.01)*** | 0.09 (0.01)*** | -0.03 (0.01) *** | -0.01 (0.01) |
| *APOE ε4* | -0.03 (0.04) | 0.01 (0.04) | -0.01 (0.04) | -0.07 (0.04) | 0.01 (0.04) | 0.02 (0.04) |
| *AKAP6* | -0.05 (0.02)* | -0.07 (0.03)* | -0.09 (0.03)*** | -0.07 (0.03)* | 0.02 (0.03) | 0.04 (0.03) |
| *MIR2113* | -0.01 (0.02) | 0.00 (0.03) | 0.00 (0.03) | 0.05 (0.03) | -0.01 (0.03) | -0.04 (0.03) |
| **Linear Rate of Change** |  |  |  |  |  |  |
| Time | -19.83 (1.71)*** | 0.58 (1.73) | 3.69 (0.91)*** | -14.79 (1.39)*** | 11.34 (2.04) *** | 13.02 (1.87) *** |
| Gender | -2.52 (1.43) | 0.35 (1.45) | 0.79 (0.76) | 1.10 (1.16) | 1.86 (1.70) | 5.01 (1.56) ** |
| Education | -0.31 (0.28) | -0.12 (0.28) | -0.01 (0.15) | -0.04 (0.23) | -0.23 (0.33) | -0.12 (0.31) |
| *APOE ε4* | -4.79 (1.47)** | -0.34 (1.50) | -0.07 (0.79) | -3.33 (1.21)** | 1.44 (1.77) | 1.44 (1.62) |
| *AKAP6* | 1.43 (1.02) | 0.79 (1.03) | 0.09 (0.54) | -0.01 (0.82) | 0.65 (1.21) | 0.05 (1.10) |
| *MIR2113* | -1.45 (1.01) | 0.34 (1.03) | -0.11 (0.54) | 0.14 (0.83) | -0.69 (1.22) | -0.60 (1.11) |
| **Quadratic Rate of Change** |  |  |  |  |  |  |
| Time<sup>2</sup> | -4.12 (1.66)*** | 0.98 (1.60) | -1.51 (0.79) | 0.31 (1.24) | -6.20 (1.94) ** | -2.52 (1.60) |
| Gender | -4.25 (1.38)** | 0.85 (1.34) | 0.39 (0.67) | 0.52 (1.03) | 1.81 (1.62) | 1.34 (1.34) |
| Education | -0.39 (0.27) | -0.33 (0.27) | -0.05 (0.13) | -0.16 (0.20) | 0.62 (0.32) | 0.57 (0.27) * |
| *APOE ε4* | 1.36 (1.42) | -1.88 (1.37) | -0.79 (0.68) | -1.48 (1.06) | 2.24 (1.66) | 2.73 (1.37) * |
| *AKAP6* | 0.44 (0.99) | -0.29 (0.96) | -0.33 (0.47) | -1.23 (0.73) | -1.81 (1.16) | -0.21 (0.95) |
| *MIR2113* | -2.43 (0.98)* | -0.82 (0.95) | -0.03 (0.47) | -0.87 (0.74) | -0.39 (1.16) | -0.17 (0.96) |
| Log Likelihood | -6601.80 | -6490.61 | -3571.27 | -5288.38 | -6981.67 | -6224.11 |
| R<sub>m</sub><sup>2</sup> | 0.19 | 0.07 | 0.22 | 0.11 | 0.07 | 0.07 |
| R<sub>c</sub><sup>2</sup> | 0.58 | 0.66 | 0.90 | 0.80 | 0.58 | 0.70 |
| Individuals | 1570 | 1570 | 1567 | 1569 | 1569 | 1568 |
| **Variance** |  |  |  |  |  |  |
| Intercept | 0.35 | 0.52 | 0.51 | 0.56 | 0.50 | 0.57 |
| Time | 83.78 | 163.22 | 56.13 | 133.11 | 199.03 | 260.50 |
| Time<sup>2</sup> | 44.15 | 62.28 | 18.78 | 43.46 | 130.81 | 74.79 |
| Residual | 0.40 | 0.32 | 0.08 | 0.18 | 0.46 | 0.30 |
| **Covariance** |  |  |  |  |  |  |
| ID x Time | -2.50 | -0.73 | -0.20 | 0.64 | 3.29 | 6.01 |
| ID x Time<sup>2</sup> | -2.69 | -0.39 | 0.76 | 0.11 | -4.57 | -1.06 |
| ID x Time x Time<sup>2</sup> | 15.91 | 45.27 | -31.19 | 65.90 | 96.04 | 108.42 |

\* $p < 0.05$ ; \*\* $p < 0.01$ .; \*\*\* $p < 0.001$

*AKAP6*-rs17522122*T was significantly associated with a lower initial status at baseline in Immediate Recall (*F*(1, 1569.2) = 3.8, *p* = 0.05), Digits Backwards test (*F*(1, 1570.0) = 5.3, *p* = 0.02), Spot-the-Word test (*F*(1, 1579.2) = 10.9, *p* = 0.0001) and Symbol Digits Modalities test (*F*(1, 1560.6) = 5.7337, *p* = 0.02) test scores. *AKAP6*-rs17522122*T was not significantly associated with baseline SRT, CRT, or linear or quadratic rate of change on any of the cognitive test scores. *MIR2113*-rs10457441*T was significantly associated with quadratic rate of change in Immediate Recall test scores showing a accelerating negative slope (*F*(1, 1217.3) = 6.5, *p* = 0.01). *MIR2113*, was not associated with any other measure of cognitive performance.

## Discussion

In this study, we investigated whether SNPs in *MIR2113* and *AKAP6*, which were previously found to be associated with cognitive performance in two large GWAS studies, were also associated with cognitive decline.

*AKAP6* was not associated with decline in any of the cognitive abilities tested. However, we did replicate the previously observed association with baseline cognitive performance, with *AKAP6*-rs17522122*T observed to be associated with worse performance in episodic memory, working memory, vocabulary and perceptual speed. In Davies *et al* 2015 & Davies *et al* 2016 *AKAP6*-rs17522122*T was associated with worse general fluid cognitive performance, verbal-numerical reasoning and improved performance in reaction time and memory. The observed differences in the direction of the effect for memory and lack of replication for reaction time between our study and Davies *et al* 2016 could be attributed to methodological differences in the cognitive tests. The cognitive tests utilized by Davies *et al* 2016 were brief bespoke, non-standard tests, with the memory test involving the recall of a single 12 item matrix that contained 6 pairs of matching symbols, with participants instructed to select the matching pairs after observing the matrix for 5 seconds in the fewest number of attempts; and reaction time assessed using 8 trials of a computerized snap game where participants were directed to push a button if two matching symbols were observed. The memory test has been shown to have low reliability across time^31^. Furthermore, both tests may be susceptible to floor/ceiling effects, where at the upper and lower limit the tests may not be able to distinguish between true differences in participant abilities^31^. Additionally, differences in participant’s demographics (i.e. age, education and comorbidities) and sample size may result in differential effects sizes, with participants in the UK Biobank been on average 5 years younger and less educated (30.5% vs 76.3% with post school qualifications) than those in PATH. Another possibility is that our study was insufficiently powered, with a dramatically smaller sample size than the UK Biobank.

*MIR2113* was associated with an accelerated rate of decline in episodic memory. We did not replicate the previous findings of the association between *MIR2113*-rs10457441*T and worse general fluid cognitive performance. It should be noted, however, that Davies *et al* 2015 used principal component analysis to derive a measure of general cognitive function from cognitive tasks testing at least three different cognitive domains. When investigating individual cognitive domains, Davies *et al* 2016 found that *MIR2113*-rs10457441*T was associated with verbal-numerical reasoning, but not with reaction time and memory. Davies *et al* 2016, suggested that the association with verbal-numerical reasoning may reflect the higher loading of verbal and reasoning abilities have on general cognitive function in comparison to memory and processing speed. As such, our results and those of Davies *et al* 2015; 2016^12,13^ suggest that *MIR2113* may be associated with general cognition rather than specific cognitive functions.

The present study has a number of strengths that allow for robust statistical modelling of the association of genetic risk factors with cognitive performance. First, this study investigated cognitive change in a randomly selected community dwelling cohort of older adults, and as such the results are likely generalizable to other population of older adults. Second, the participants were followed for 12 years with four follow-up interviews assessing four separate cognitive domains, allowing for the examination of non-linear declines across a broad spectrum of cognitive abilities. Finally, the narrow age range of the cohort reduces the potential impact of the confounding effects of age. Despite these strengths, the results from this study should be interpreted with consideration to some study limitations. First, the 60+ cohort is better educated than the population from which was drawn. As higher education is associated with a reduced rate of decline this may limit our ability to observe an association with cognitive performance. Second, only the most strongly associated SNP identified by the CHARGE consortium at each locus was genotyped. These SNPs are unlikely to be the causal variants, but are expected to be in linkage disequilibrium with other genetic variation that causes the observed associations with cognitive function. Further, ‘gene’ based analyses that aggregate the effects of multiple SNPs may further elucidate the role of these loci in cognitive performance.

In conclusion, we have shown that *AKAP6* does not influence the rate of cognitive decline in a population of healthy older adults. However, *AKAP6* is associated with baseline cognitive performance across a broad spectrum of cognitive domains. In contrast, *MIR2113* was associated with an accelerated decline in episodic memory, however, we did not observe an association with baseline cognitive performance. These results extend upon those reported by Davies *et al* ^12,13^, further elucidating the role these of loci in cognitive performance.

## Acknowledgments

We thank the investigators in the PATH study: Peter Butterworth, Andrew Mackinnon, Anthony Jorm, Bryan Rodgers, Helen Christensen, Patricia Jacomb and Karen Maxwell. The study was supported by the National Health and Medical Research Council (NHMRC) grants 973302, 179805, 1002160 and 1002160 and the NHMRC Dementia Collaborative Research Centre Early Diagnosis and Prevention. DD is funded by NHMRC Project Grant No. 1043256. KJA is funded by NHMRC Research Fellowship No. 1002560.

